# Deregulation of TGF-β1 signaling induces glycolysis by chromatin remodeling in pathogenic TH17 cells

**DOI:** 10.1101/218545

**Authors:** Xiang Yu, Li Wang, Zhijun Han, Chao Yao, Rong Qiu, Yange Cui, Dai Dai, Wenfei Jin, Nan Shen

## Abstract

It is well known that some pathogenic cells have enhanced glycolysis, the regulatory network leading to increased glycolysis are not well characterized. Here, we show that pathogenic T_H_17 cells specifically upregulate glycolytic pathway genes compared to homeostatic T_H_17 cells. Bioenergetic assay and metabolomics analyses indicate that pathogenic T_H_17 cells are highly glycolytic compared to nonpathogenic T_H_17 cells. Chromatin landscape analyses demonstrate T_H_17 cells in vivo show distinct chromatin states, and pathogenic T_H_17 cells show enhanced chromatin accessibility at glycolytic genes with NF-kB binding sites. Mechanistic studies reveal that TGF-β1 signaling induces T_H_17 cell chromatin remodeling and represses c-Rel-mediated glycolysis. A *miR-21-Peli1*-c-Rel loop was further identified to be essential for glycolysis of pathogenic T_H_17 cells. These findings extend our understanding of the regulation T_H_17 cell glycolysis in vivo and provide insights for future therapeutic intervention to T_H_17 cell mediated autoimmune diseases.

## Introduction

T_H_17 cells, which have been characterized as a subset of effector CD4^+^ helper T cells that produce IL-17A (Littman and Rudensky, 2010), mediate barrier tissue integrity, mucosal defence, and also contribute to the development of multiple autoimmune diseases. At steady-state condition, T_H_17 cells producing IL-10 are mainly found at sites of intestinal lamina propria (Ivanov et al., 2009; Sano et al., 2015). Under autoimmune or pathogen infection condition, T_H_17 cells acquire the capability to produce IFNγ or GM-CSF, and are often seen at sites of inflamed loci (Atarashi et al., 2015; Stockinger and Omenetti, 2017). T_H_17 cells also could be differentiated in vitro by a combination of cytokines, either by TGF-β1 plus IL-6 (T_H_17 (β), nonpathogenic) or by IL-6 plus IL-1β and IL-23 (T_H_17 (23), pathogenic) (Gaublomme et al., 2015; Ghoreschi et al., 2010).

Upon antigen stimulation, lymphocytes undergo extensive clonal expansion and differentiation for immune defense or tolerance (Boothby and Rickert, 2017; Buck et al., 2015; Man and Kallies, 2015; Pollizzi and Powell, 2014). Activated lymphocytes are highly glycolytic and demonstrate a striking increase in cell content and nutrients uptake. c-Myc, c-Rel and mTORc1-dependent glycolytic activity is crucial for effective T and B cell immune response (Heise et al., 2014; Karmaus et al., 2019; Verbist et al., 2016; Wang et al., 2011), whereas Foxo1 and Foxp3 proteins function as key metabolic repressors that limit glycolytic activity in Treg cells (Angelin et al., 2017; Cho et al., 2016; Wilhelm et al., 2016). Recently it is reported that by restricting glucose availability, tumor cells limit aerobic glycolysis and effector function of tumor-infiltrating T cells (Chang et al., 2015; Ho et al., 2015). However, metabolic states of T_H_17 cells in vivo and its regulation under homeostatic condition is incompletely understood.

Tight communication between epigenetic remodeling and metabolic regulation for cell plasticity is beginning to be understood (Etchegaray and Mostoslavsky, 2016; Rajbhandari et al., 2018). Assay for transposase-accessible chromatin technology (ATAC-seq) was developed to study global chromatin dynamics by using relatively low cell number (Buenrostro et al., 2013). It was successfully used to study chromatin dynamics of Treg cells under inflammatory condition (van der Veeken et al., 2016), development of plasmacytoid DC (Ceribelli et al., 2016) and tumor-specific CD8^+^ T cells (Philip et al., 2017). However, chromatin architecture of T_H_17 cells in vivo is unknown, and signaling pathways that induce T_H_17 cell chromatin remodeling are also incompletely understood.

TGF-β1 is a cytokine with immunosuppressive properties and its loss is associated with fatal autoimmune pathologies (Fontenot et al., 2003; Gorelik and Flavell, 2000; Shull et al., 1992). TGF-β1 is secreted by multiple cell types, including regulatory T cells, fibroblasts, and epithelial cells. It signals through a receptor complex of TGF-βRII and TGF-βRI to trigger the phosphorylation of Smad proteins, which then act to induce and repress specific target genes (Tu et al., 2018). TGF-β1, at a lower concentration, promotes nonpathogenic T_H_17 cell differentiation in vitro (Ghoreschi et al., 2010; McGeachy et al., 2007), however, at a relatively higher concentration, it readily induces Treg cells (Li and Flavell, 2008). However, the precise mechanism by which TGF-β1 regulates T_H_17 cell function remains poorly understood.

In this study, first by global transcriptional analyses we demonstrate that homeostatic T_H_17 cells derived in vivo show repressed glycolytic pathway gene expression. Further by bioenergetic and metabolomics study we demonstrate that in vitro derived nonpathogenic T_H_17 (β) cells show reduced glycolytic activity compared to pathogenic T_H_17 (23) cells. Chromatin landscape analyses indicate that T_H_17 cells in vivo show distinct chromatin states. Pathogenic T_H_17 cells-specific open chromatin region are enriched for motifs of NF-kB family transcription factors, whereas homeostatic T_H_17 cells show repressed chromatin accessibility at glycolytic genes with NF-kB binding sites. Mechanistic studies reveal TGF-β1 signaling induces T_H_17 cell chromatin remodeling and represses c-Rel-mediated glycolysis to maintain metabolic quiescence of nonpathogenic T_H_17 cells. A *miR-21-Peli1*-c-Rel loop was further identified to be essential for glycolysis of pathogenic T_H_17 cells.

## Results

### Homeostatic T_H_17 cells show repressed glycolysis pathway gene expression

To gain insight of the metabolic states of T_H_17 cells in vivo, we sorted GFP^+^ T_H_17 cells from homeostatic ileum and inflamed CNS of EAE mice (Figure S1A). Then we sent ileum and CNS-infiltrated T_H_17 cells for RNA-seq analyses. We found 1415 upregulated genes and 1249 downregulated genes in CNS-infiltrated T_H_17 cells compared to ileum T_H_17 cells (Figure S1B-C). GO and Pathway analyses indicate that several immune pathways were activated in CNS-infiltrated T_H_17 cells as reported (Gaublomme et al., 2015), interestingly, we found multiple metabolic pathways were enriched (Figure 1A-F, Figure S1D). We found homeostatic ileum T_H_17 cells express great more *Foxp1 gene* (Figure 1A), which is critical for maintenance of the metabolic quiescence of naive T cells (Feng et al., 2011; Stephen et al., 2014). Ileum homeostatic T_H_17 cells also express several transcription factors with repressor activity *c-Maf* and *Foxo1*, and Foxo1 was a repressor of glucose metabolism (Wilhelm et al., 2016) (Figure 1A), which is consistent with our finding that CNS-infiltrated T_H_17 cells express higher level of glycolysis pathway genes (Figure 1B-D, Figure S1D), whereas ileum T_H_17 cells show higher enrichment of glutaminolysis pathway genes. Taken together, these data suggest that T_H_17 cells in vivo have differential expression of key metabolic pathway genes, with ileum homeostatic T_H_17 cells show repressed glycolysis pathway gene activation.

**Figure 1.**
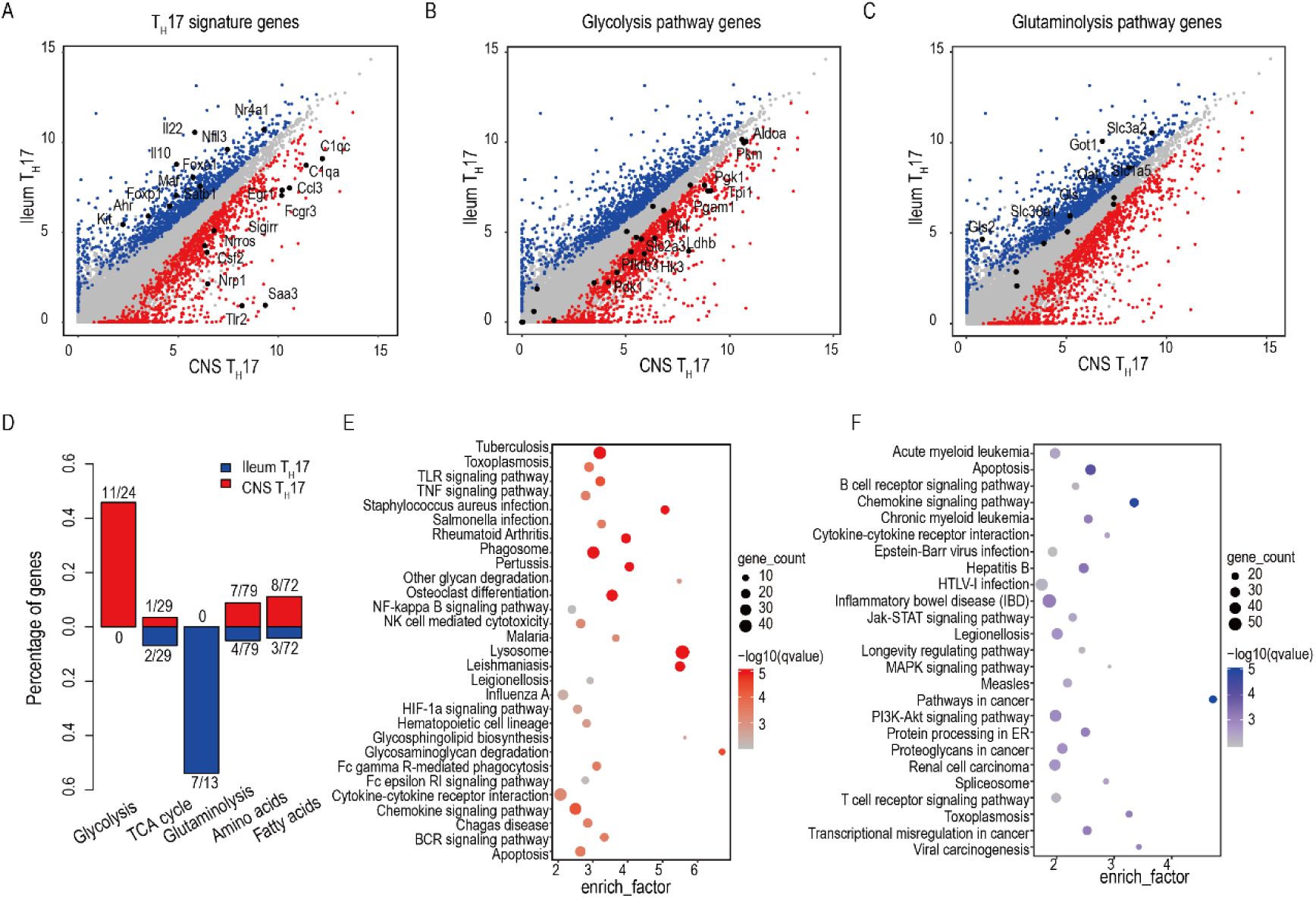
Homeostatic T_H_17 cells show repressed glycolysis pathway gene expression (A) Scatter plot shows differential expressed genes between CNS T_H_17 (red) and Ileum T_H_17 (blue) with differential T_H_17 signature genes highlighted. DEGs are filtered as FDR < 0.05 and fold change ≥ 1.5 from DESeq2. (B) Scatter plot shows differential expressed genes between CNS T_H_17 (red) and Ileum T_H_17 (blue) with glycolysis pathway genes highlighted and differential genes labeled. (C) Scatter plot shows differential expressed genes between CNS T_H_17 (red) and Ileum T_H_17 (blue) with glutaminolysis pathway genes highlighted and differential genes labeled. (D) CNS T_H_17 and Ileum T_H_17 cells depend on distinct metabolic pathways. Numbers indicate the differential expressed genes and total genes for each pathway. (E) KEGG enrichment for CNS T_H_17 cell highly expressed genes. (F) KEGG enrichment for Ileum T_H_17 cell highly expressed genes.

### Nonpathogenic T_H_17 cells show reduced glycolytic activity to pathogenic T_H_17 cells

To study the metabolic activity of T_H_17 cells more deeply, we induced T_H_17 cells in vitro by a combination of cytokines TGF-β1, IL-6 (T_H_17 (β)) or IL-6, IL-1β and IL-23 (T_H_17 (23)). Interestingly, we consistently observed that T_H_17 (23) cells show increased cell size compared to T_H_17 (β) cells (Figure S2A-B). We further tested the glycolytic activity of T_H_17 (β) and T_H_17 (23) cells by analyzing key metabolic enzyme gene expression, we found T_H_17 (23) cells express higher glycolytic pathway genes than T_H_17 (β) cells, T_H_17 (23) cells express great more *Slc2a1/3, Hk2, Tpi1, Aldoa, Eno1, Pkm and Ldha* (Figure 2A, Figure S2C). We further checked the glycolytic activity of T_H_17 (β) and T_H_17 (23) cells by glycolysis stress test, both basal and maximal glycolytic capability were much higher in pathogenic T_H_17 (23) cells (Figure 2B).

**Figure 2.**
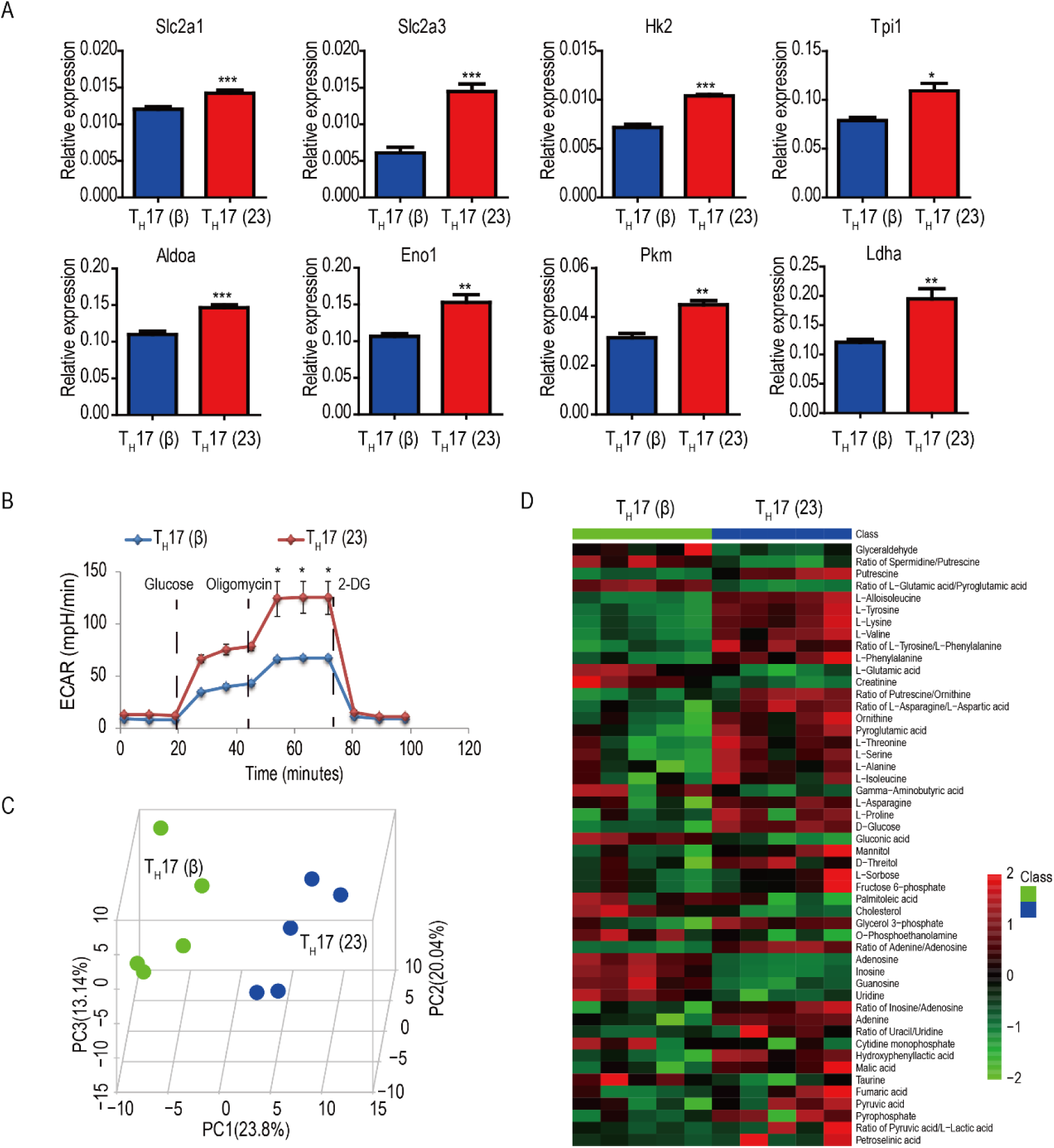
Nonpathogenic T_H_17 cells show reduced glycolytic activity to pathogenic T_H_17 cells (A) RT-PCR analysis of key glycolytic pathway genes in T_H_17(β) and T_H_17(23) cells differentiated in vitro for 48 hs. (B) Extracellular acidification rate (ECAR) of T_H_17 (β) and T_H_17 (23) cells differentiated for 96 hs assessed by a glycolytic stress test. (C) PCA analysis of identified metabolites of T_H_17 (β) and T_H_17 (23) cells differentiated in vitro for 48 hs by metabolomics. (D) Metabolomics analysis of T_H_17 (β) and T_H_17 (23) cells differentiated in vitro for 48 hs, the top differentially observed metabolites are shown in heat map. *P < 0.05 (unpaired t-test). Data are from one experiment representative of three independent experiments (A).

We next subjected T_H_17 (β) and T_H_17 (23) cells for high-resolution metabolomics analyses (Chen et al., 2014; Chen et al., 2016). Pathogenic T_H_17 (23) cells were metabolically distinct from T_H_17 (β) cells (Figure 2C, Figure S2D), and showed higher level intermediates of carbohydrates, amino acids and organic acids, while T_H_17 (β) cells showed higher level intermediates of nucleotides and fatty acids. Pathway analyses of altered metabolites showed that amino acid, amino sugar metabolism, and glycolytic intermediates trended to be higher in T_H_17 (23) cells (Figure 2D, Table S1). Taken together, these data suggest that T_H_17 cells in vitro have distinct metabolic states, nonpathogenic T_H_17 (β) cells are less glycolytic compared to pathogenic T_H_17 (23) cells.

### Homeostatic T_H_17 cells show repressed chromatin accessibility at glycolytic genes

To dissect possible regulation mechanism for T_H_17 cell metabolic reprogramming in vivo, we performed ATAC-seq to assess chromatin landscape genome-wide of T_H_17 cells derived in vivo (Figure S3A-C). We found that T_H_17 cells had discrete chromatin states in vivo (Figure 3A-B), CNS infiltrating T_H_17 cells showed great differential global chromatin accessibility compared to ileum homeostatic T_H_17 cells. Transcriptional factor motif analysis of top 1000 enriched peaks indicated that NF-kB family members (RelA, c-Rel) were repressed in homeostatic T_H_17 cells (Figure 3C). Interestingly, the regulatory region of key T_H_17 cell-related transcription factors c-*Maf* and *Ahr* was with enhanced chromatin accessibility in homeostatic T_H_17 cells, both c-Maf and Ahr were shown to be critical regulators of T_H_17 cells (Figure S3E). The regulatory region of metabolic repressor Foxo1 was with reduced chromatin accessibility in CNS-infiltrated T_H_17 cells, which suggests that CNS-infiltrated T_H_17 cells were relieved from Foxo1-mediated metabolic repression (Figure S3E). A group of genes related to metabolic pathways and solute carrier family members were also with repressed chromatin accessibility in steady-state ileum T_H_17 cells, which was consistent with the gene expression data (Figure 3D, Figure S3F). Taken together, these data suggest that metabolic reprogramming of T_H_17 cells in vivo were controlled at the epigenetic level, homeostatic T_H_17 cells show repressed chromatin accessibility at glycolytic genes.

**Figure 3.**
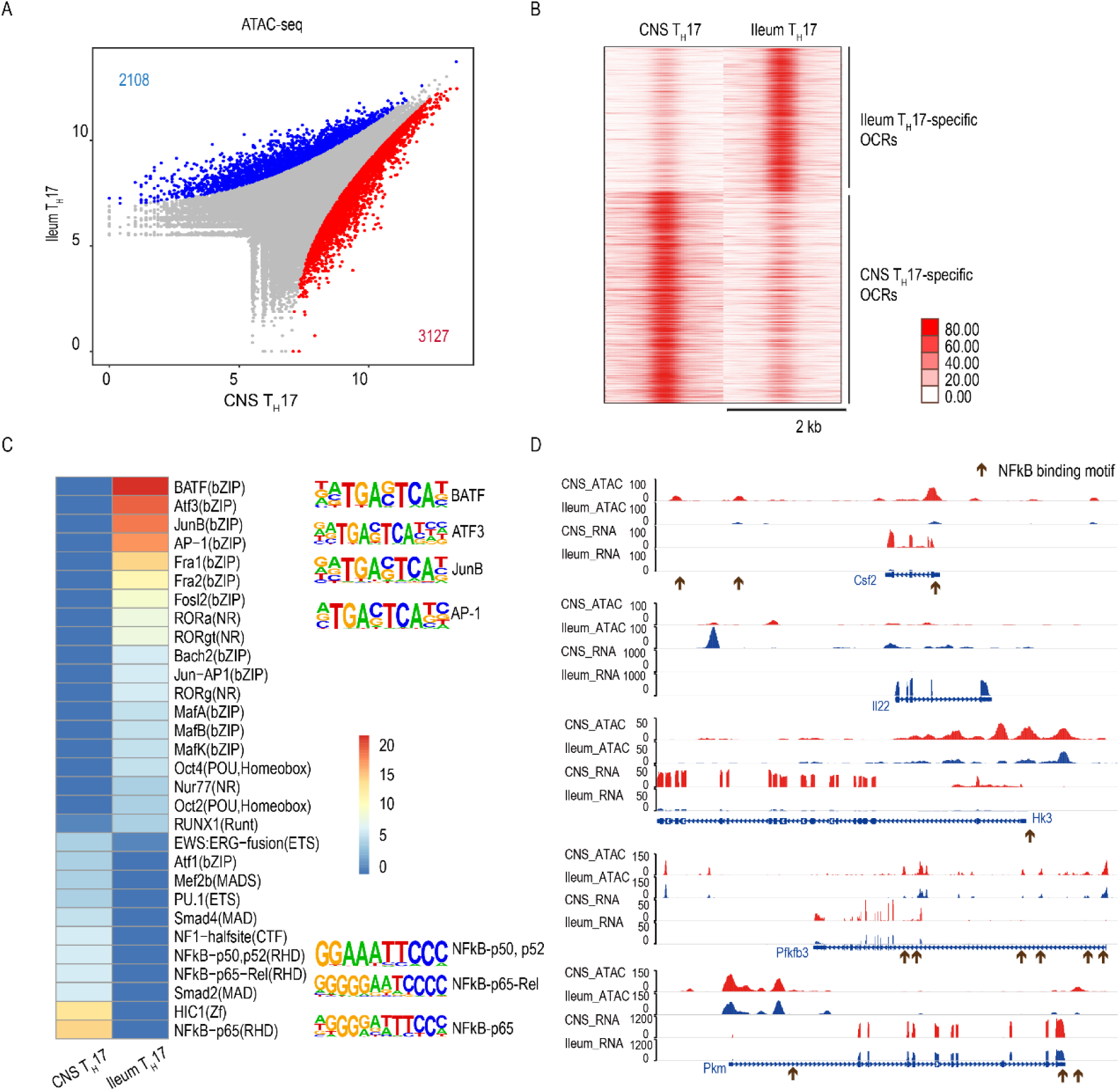
Homeostatic T_H_17 cells show repressed chromatin accessibility at glycolytic genes (A) Scatter plot for differential opened/closed regions (OCRs) between CNS T_H_17 (red) and Ileum T_H_17 cells (blue) with differential numbers labeled. (B) ATAC-seq reads density heatmap around the OCRs between CNS T_H_17 and Ileum T_H_17 cells. (C) Enriched transcription factor motifs in top 1000 differential OCRs between CNS T_H_17 cells and Ileum T_H_17 cells, color bar is −log10(P value). Right panel shows the identified motif sequences for key regulators. (D) Differential OCRs signals with corresponding gene expression levels for key T_H_17 signature genes and metabolic genes in CNS T_H_17 and Ileum T_H_17 cells. Arrows represent the NF-kB binding motif sites.

### Deregulation of TGF-β1 signaling induces T_H_17 cell chromatin remodeling at glycolytic genes

To define the regulation mechanism for T_H_17 cell chromatin remodeling, we differentiated T_H_17 cells in vitro either in the presence (T_H_17 (β)) or absence (T_H_17 (23)) of TGF-β1 and sorted out GFP^+^ T_H_17 cells, then we performed ATAC-seq to assess chromatin landscape genome-wide (Figure S4A-B). We found T_H_17 cells derived in vitro had discrete chromatin states (Figure 4A), T_H_17 (23) cells showed great differential global chromatin accessibility compared to T_H_17 (β) cells (Figure 4B). Transcriptional factor motif analysis of enriched peaks indicated that NF-kB family members were repressed in nonpathogenic T_H_17 (β) cells derived in the presence of TGF-β1 signaling (Figure 4C), which is consistent with the reports that TGF-β1 signaling represses c-Rel activity during lymphocytes activation (Arsura et al., 1996). A group of genes related to glycolytic pathway were also with decreased chromatin accessibility in T_H_17 cells derived in the presence of TGF-β1 signaling, which was also consistent with their relative expression level (Figure 4D, Figure 2A). Taken together, these data suggest that TGF-β1 signaling represses T_H_17 cell chromatin accessibility at glycolytic genes for metabolic reprogramming.

**Figure 4.**
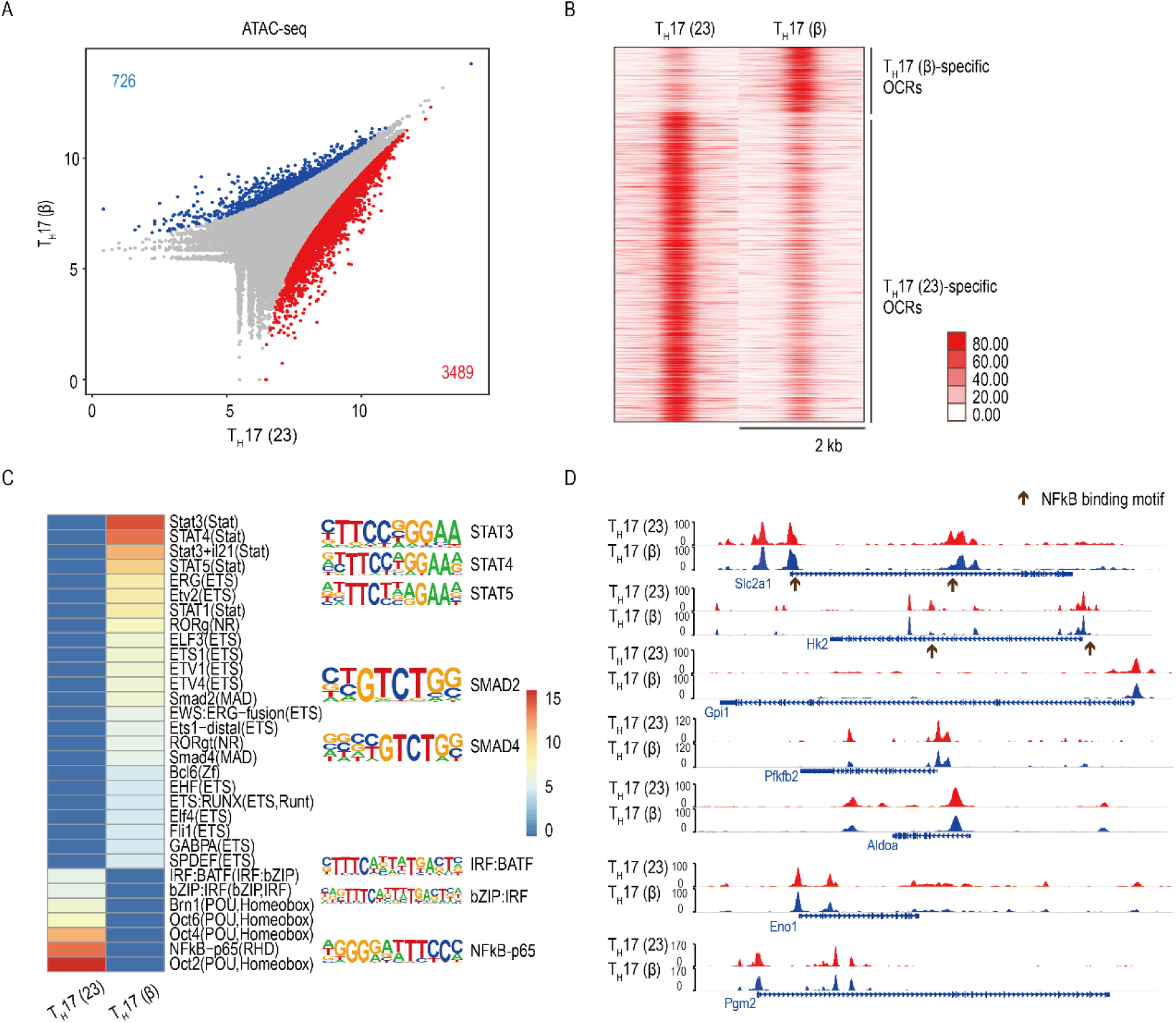
Deregulation of TGF-β1 signaling induces T_H_17 cell chromatin remodeling at glycolytic genes (A) Scatter plot for differential opened/closed regions (OCRs) between T_H_17 (23) (red) and T_H_17 (β) cells (blue) with differential numbers labeled. (B) ATAC-seq reads density heatmap around the OCRs between T_H_17 (23) and T_H_17 (β) cells. (C) Enriched transcription factor motifs in top 1000 differential OCRs between T_H_17 (23) cells and T_H_17 (β) cells, color bar is −log10 (P value). Right panel shows the identified motif sequences for key regulators. (D) Differential OCRs signals for key glycolytic genes in T_H_17 (23) and T_H_17 (β) cells. Arrows represent the NF-kB binding motif sites.

### TGF-β1 signaling represses c-Rel-mediated glycolysis pathway gene activation in nonpathogenic T_H_17 cells

Our above data suggested that genes with NF-kB family transcription factors motif were repressed in homeostatic ileum T_H_17 cells and in vitro-derived nonpathogenic T_H_17 cells at the chromatin level. To further explore the role of TGF-β1 signaling on NF-kB family transcription factors and its effect on T_H_17 cell glycolysis, we differentiated T_H_17 cells in vitro under nonpathogenic T_H_17 (β) cell condition with variable concentration of mouse recombinant TGF-β1, we found that TGF-β1 signaling inhibited c-Rel expression in a dose-dependent manner, while RelA expression was unchanged at the protein level (Figure 5A). We further checked c-Rel and RelA protein level at in vitro-derived nonpathogenic T_H_17 (β) and pathogenic T_H_17 (23) cells, we found pathogenic T_H_17 (23) cells expressed specifically more c-Rel protein than nonpathogenic T_H_17 (β) cells (Figure 5B). To study whether c-Rel was essential for glycolysis pathway gene activation of pathogenic T_H_17 cells, we took advantage of a c-Rel specific inhibitor pentoxifylline (PTXF) (Grinberg-Bleyer et al., 2017), which was approved by FDA for clinical use for a variety of diseases. Inhibition of c-Rel reduced the expression key glycolytic pathway gene expression in activated pathogenic T_H_17 (23) cells (Figure 5C-D), including *Slc2a3*, *Aldoa*, *Tpi1*, *Eno1* and *Ldha*. Next, we asked whether overexpression of c-Rel in nonpathogenic T_H_17 (β) cells could promote pathogenic T_H_17 cell signature genes and glycolytic pathway gene expression, indeed enforced expression of c-Rel in T_H_17 (β) cells promoted key pathogenic T_H_17 cell signature gene expression (Figure 5E, Figure S5A-B), including *IL17a*, *IL17f*, *Csf2*, *IL2* and *IL23r*. And we found enforced expression of c-Rel promoted selected glycolysis pathway gene expression with a level of 20% to 50% upregulation (*Hk1*, *Hk2*, *Slc2a1* and *Slc2a3*) (Figure 5F), though less pronounced to the level of upregulation of pathogenic T_H_17 signature genes. We further checked key glycolytic enzymes at the protein level, and we found overexpression of c-Rel elevated Slc2a1 (Glut1) and HK2 (Figure 5G). Together, these data suggest that TGF-β1 signaling represses c-Rel-mediated glycolysis of pathogenic T_H_17 cells.

**Figure 5.**
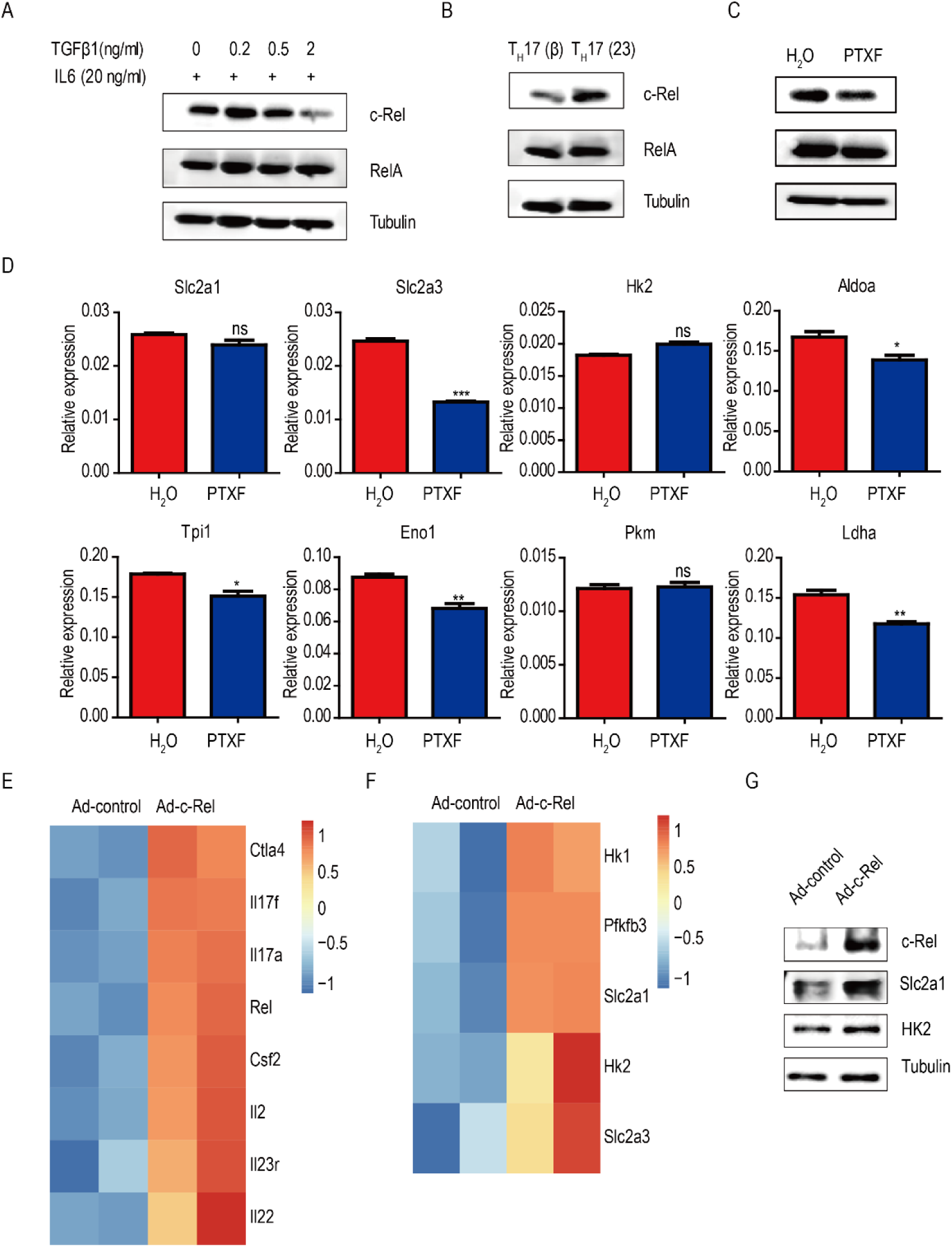
TGF-β1 signaling represses c-Rel-mediated glycolysis pathway gene activation in nonpathogenic T_H_17 cells (A) Western blot analysis of c-Rel and RelA from T_H_17 cells differentiated under indicated condition. (B) Western blot analysis of c-Rel and RelA from T_H_17 cells differentiated under indicated condition. (C) Western blot analysis of c-Rel, RelA from T_H_17 cells differentiated under indicated condition. (D) RT-PCR analysis of key glycolytic pathway genes in T_H_17(23) cells differentiated in vitro for 48 hs, treated with H_2_O or 500 µg/ml PTXF for 16-18 hs. (E) Heat map of selected genes of interest in T_H_17 cell pathogenicity (FDR < 0.05, fc ≥ 1.5). (F) Heat map of selected genes of interest in glycolysis pathway (FDR < 0.05, fc ≥ 1.2). (G) Western blot analysis of c-Rel, Slc2a1 and HK2 from T_H_17 cells differentiated under indicated condition. *P < 0.05 (unpaired t-test). Data are from one experiment representative of three independent experiments (A, B, C, D and G).

### A miR-21-Peli1-c-Rel loop promotes pathogenic T_H_17 cell glycolysis

Specific microRNAs are appreciated as key metabolic regulators for immune cell function, to identify microRNAs activated by NF-kB transcription factors that are essential for glycolysis of pathogenic T_H_17 cells, we checked chromatin accessibility of the regulatory region of microRNAs with NF-kB binding sites and their expression in T_H_17 cells derived in vivo. We found the regulatory region of a group of microRNAs show great more chromatin accessibility in CNS pathogenic T_H_17 cells, which include *miR-21, miR-23a cluster, miR-146a* (Figure 6A-B), while miR-21 and miR-146a were the two with the highest abundance in CNS T_H_17 cells.

**Figure 6.**
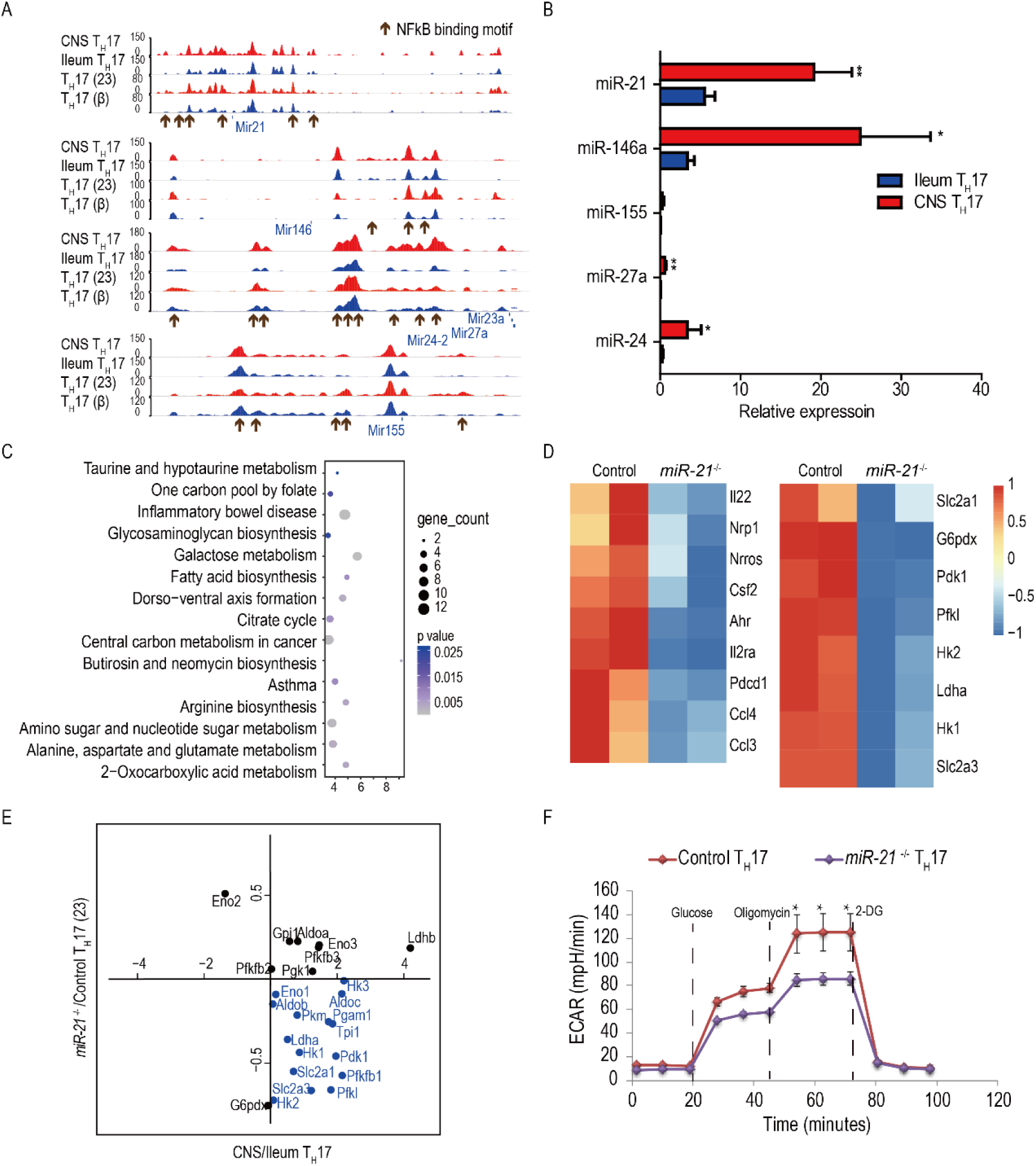
A miR-21-Peli1-c-Rel loop promotes pathogenic T_H_17 cell glycolysis (A) ATAC-seq signal profile across selected microRNA genome loci of ileum homeostatic and CNS-infiltrated pathogenic T_H_17 cells. (B) Relative expression of microRNA in ileum homeostatic and CNS-infiltrated pathogenic T_H_17 cells (n=3). (C) KEGG enrichment for genes downregulated in *miR-21*^−/−^ T_H_17 cells. (D) Heat map of selected genes of interest in T_H_17 cell pathogenicity (left) and in glycolytic pathway (right). (E) RNA-seq data of glycolytic pathway genes from *miR-21*^−/−^/control T_H_17 cell ratio plotted as a function of RNA-seq data from CNS/Ileum T_H_17 cell ratio. (F) Extracellular acidification rate (ECAR) of T_H_17 (23) cells differentiated from control and *miR-21*^−/−^ mice for 96 hs assessed by a glycolysis stress test.

Our group previously demonstrated that miR-21 was highly upregulated in inflammatory lesions of multiple human autoimmune diseases (Zhu et al., 2012). To better understand the general functional features of miR-21-deficient T_H_17 cells, we differentiated naïve T cells from control and *miR-21*^−/−^ mice in vitro under pathogenic T_H_17 (23) condition, RNA was extracted and subjected to RNA-seq. In the absence of miR-21, T_H_17 cells showed decreased expression of T_H_17 cell signature genes, which include lineage-specific transcription factors, cytokines, and cytokine receptors (Figure 6C-D). Remarkably, many of the genes involved in glycolytic or related metabolic pathways were highly downregulated, including key transporters for glucose intake, *Slc2a1/3*, and rate-limiting enzymes, *Hk1/2*, *Pfkl*, *Pgm2*, *Ldha* and *Pdk1* (Figure 6D). Interestingly, many genes of the glycolytic pathway that are highly expressed in CNS-infiltrated pathogenic T_H_17 cells are generally downregulated in *miR-21*^−/−^ T_H_17 cells (Figure 6E). We further checked the glycolytic activity of control and *miR-21*^−/−^ T_H_17 cells by glycolysis stress test, both basal and maximal glycolytic capability were decreased in *miR-21*^−/−^ T_H_17 cells (Figure 6F). Taken together, these data suggest that miR-21 promotes glucose metabolism of pathogenic T_H_17 cells.

To further investigate the precise mechanism underlying how miR-21 promotes pathogenic T_H_17 cell glucose metabolism, we screened direct targets of miR-21 by combining target prediction with Ago2 HITS-CLIP (Loeb et al., 2012), TargetScan and RNA-seq data to identify overlapping gene transcripts. Using a combination of these three approaches, we identified 13 common putative targets (Figure S6A). Among them, *Peli1* has been reported to function as a negative regulator of CD4^+^ T cell activation by ubiquitination of c-Rel (Chang et al., 2011), therefore we chose *Peli1* for further analysis. By in vitro RIP assay, we found Ago2-immunoprecipitated RNAs had significantly reduced Peli1 mRNA level in *miR-21*^−/−^ T_H_17 (23) cells (Figure S6B-C), which suggests that Peli1 was a functional target of miR-21 in pathogenic T_H_17 cells. We further validated these at the protein level in purified GFP^+^ T_H_17 cells, we found that *miR-21*^−/−^ T_H_17 cells have elevated level of Peli1 protein, whereas the protein level of c-Rel was reduced in *miR-21*^−/−^ T_H_17 cells (Figure S6D). Taken together, these data indicate that a *miR-21-Peli1*-c-Rel loop promotes glycolysis of pathogenic T_H_17 cells.

## Discussion

T_H_17 cells exhibit great heterogeneity in vivo, the basic energy requirement of T_H_17 cells for their regulatory or inflammatory function under homeostatic or autoimmune condition was not examined. And how T_H_17 cells adapt to diverse tissue microenvironment with distinct nutrients availability and regulation of their metabolic reprogramming in vivo were not known.

In the present study, by metabolomics analyses of in vitro derived T_H_17 cells, we found that T_H_17 cells respond to different stimulations with distinct metabolic activity. And by global transcriptome analyses of in vivo derived T_H_17 cells, we first showed T_H_17 cells adapt to tissue microenvironment with discrete metabolic pathway gene activation in vivo.

To dissect the regulation of T_H_17 cell metabolic states in vivo, we performed genome-wide chromatin landscape profiling and found that T_H_17 cells had discrete chromatin states in vivo. The regulatory region of key T_H_17 cell-related transcription factors, metabolic regulators, metabolism-related genes and microRNAs were with differential chromatin accessibility between homeostatic ileum T_H_17 cells and CNS infiltrating T_H_17 cells. Our data suggests that T_H_17 cells in vivo adapt to distinct microenvironment with discrete epigenetic and metabolic states, and metabolic regulators and metabolism-related genes were regulated at the epigenetic level for their functional requirements. TGF-β1 signaling was further shown to be crucial for remodeling of T_H_17 cell chromatin states in vitro, c-Rel was identified as an essential metabolic regulator for pathogenic T_H_17 cells.

Interestingly, we found the conserved regulatory region of a group of small microRNAs were with differential chromatin accessibility of T_H_17 cells derived in vivo, which could have contributed to the regulation of discrete metabolic states of T_H_17 cells in vivo for the fine-tune property of microRNAs. The regulatory region of miR-21 was with increased chromatin accessibility in CNS infiltrating T_H_17 cells. Although miR-21 was shown previously to be important in T_H_17 cell generation (Murugaiyan et al., 2015), its exact action has not been well understood. By global transcriptomics analyses, we demonstrate that miR-21^−/−^ T_H_17 cells express less T_H_17 cell signature genes and key metabolism pathway genes. These data suggest that with increased chromatin openness of its regulatory region, miR-21 regulates metabolic reprogramming of T_H_17 cells under autoimmune inflammation.

T_H_17 cells mediate pathogenic role in multiple autoimmune diseases. Though anti-IL-17A treatment received great efficacy in psoriasis, it badly failed for IBD patients. Understanding metabolic reprogramming of T_H_17 cells in vivo may therefore provide more defined therapeutic intervention to T_H_17 cell mediated autoimmune diseases and insights into T_H_17 cell mediated host defence. The microenvironment of homeostatic mucosal surfaces and various inflammatory sites could have great differential effects on chromatin states of T_H_17 cells in vivo, which in turn could facilitate their discrete metabolic reprogramming. Our study so far dissected the metabolic states and chromatin states of T_H_17 cells in vivo and identified c-Rel as a critical metabolic regulator of pathogenic T_H_17 cells.

## STAR Methods

### Key Resources Table

| REAGENT or RESOURCE | SOURCE | IDENTIFIER |
| --- | --- | --- |
| Antibodies |  |  |
| Anti-IFNγ functional | ThermoFisher | Catalog # 16-7311-38 |
| Anti-IL4 functional | ThermoFisher | Catalog # 16-7041-95 |
| Anti-CD3 functional | ThermoFisher | Catalog # 16-0038-85 |
| Anti-CD28 functional | ThermoFisher | Catalog # 16-0289-85 |
| Foxo1 | Cell Signaling | Catalog # 2880S |
| c-Myc | Cell Signaling | Catalog # 13987S |
| Phospho-S6 | Cell Signaling | Catalog # 4856S |
| HK2 | Cell Signaling | Catalog # 2867S |
| Slc2a1 | Abcam | Catalog # ab115730 |
| Peli1 | Cell Signaling | Catalog # 31474S |
| c-Rel | Cell Signaling | Catalog # 12707S |
| GAPDH | Cell Signaling | Catalog # 3683S |
| Tubulin | Cell Signaling | Catalog # 2146S |
| Anti-Mouse/Rat Foxp3 APC | eBioscience | Catalog # 17-5773-82 |
| Anti-Mouse CD3e PE-Cyanine5.5 | eBioscience | Catalog # 35-0031-82 |
| PE Rat Anti-Mouse CD4 Clone RM4-5 | BD Biosciences | Catalog # 553048 |
| APC-Cy<sup>TM</sup>7 Rat Anti-Mouse CD45 | BD Biosciences | Catalog # 557659 |
| PE Mouse anti-Mouse RORγt | BD Biosciences | Catalog # 562607 |
| PerCP-Cy<sup>TM</sup>5.5 Rat Anti-Mouse IL-17A | BD Biosciences | Catalog # 560666 |
| APC Rat Anti-Mouse IFN-γ | BD Biosciences | Catalog # 554413 |
| BV421 Rat Anti-Mouse CD4 | BD Biosciences | Catalog # 562891 |
| PE Rat Anti-Mouse GM-CSF | BD Biosciences | Catalog # 554406 |
| Chemicals, Peptides, and Recombinant Proteins |  |  |
| Corning® Cell-Tak<sup>TM</sup> Cell and Tissue Adhesive | Corning | Catalog # 352441 |
| Recombinant Mouse IL-6 Protein | R & D systems | Catalog # 406-ML-025 |
| Recombinant Mouse IL-1 beta/IL-1F2 Protein | R & D systems | Catalog # 401-ML-025 |
| Recombinant Mouse IL-23 Protein | R & D systems | Catalog # 1887-ML-010 |
| Recombinant Mouse TGF-beta 1 Protein | R & D systems | Catalog # 7666-MB-005 |
| Percoll | Sigma | Catalog # 17-0891-01 |
| Pertussis toxin | Sigma | Catalog # P7208 |
| Pentoxifylline | Sigma | Catalog # P1784 |
| Adjuvant, Incomplete (Freund), 10 mL | BD Difco | Catalog # 263910 |
| M. Tuberculosis H37 Ra, Desiccated, 100 mg | BD Difco | Catalog # 231141 |
| Myelin Oligodendrocyte Glycoprotein (MOG 35-55) | Chinapeptides | Catalog # 163913-87-9 |
| **Critical Commercial Assays** |  |  |
| LIVE/DEAD™ Fixable Violet Dead Cell Stain Kit, for 405 nm excitation | ThermoFisher | Catalog # L34955 |
| Naive CD4+ T Cell Isolation Kit | Miltenyi Biotec | Catalog # 130-104-453 |
| Seahorse XF Glycolysis Stress Test Kit | Agilent technologies | Catalog # 103020-100 |
| miScript Single Cell qPCR Kit (24) | Qiagen | Catalog # 331053 |
| Magna RIP™ RNA-Binding Protein Immunoprecipitation Kit | Merck Millipore | Catalog # 17-700 |
| Nextera® DNA Sample Preparation Kit (24 Samples) | illumina | Catalog # FC-121-1030 |
| miRNeasy Micro Kit | Qiagen | Catalog # 217084 |
| **Deposited Data** |  |  |
| All sequencing data | This paper | GEO: GSE127768 |
| **Experimental Models: Organisms/Strains** |  |  |
| IL17A-eGFP | Biocytogen | N/A |
| *miR-21* -/- | Shanghai Model Organisms | Stock No: NM-KO-00159 |
| *miR-21* f/+ | Shanghai Model Organisms | Stock No: NM-CKO-00051 |
| **Oligonucleotides** |  |  |
| has-miR-21 | ABI | Catalog # 000397 |
| U6 snRNA | ABI | Catalog # 001973 |
| Primers for qPCR see Table S2 |  |  |
| **Software and Algorithms** |  |  |
| Flowjo version V10 | Flowjo | https://www.flowjo.com |
| GraphPad Prism v6.0 | GraphPad Software | https://www.graphpad.com |
| Agilent Seahorse Wave 2.4 | Agilent technologies | [http://www.agilent.com/en-us/products/cell-analysis-(seahorse)/seahorse-wave-software](http://www.agilent.com/en-us/products/cell-analysis-%28seahorse%29/seahorse-wave-software) |

### Contact for Reagent and Resource Sharing

Further information and requests for resources and reagents should be directed to and will be fulfilled by the Lead Contact, Nan Shen

### Experimental Model and Subject Details

#### Mice

*MiR-21*^−/−^ and *miR-21* ^f/+^ mice on the C57B6 background were from Shanghai Model Organisms (China). IL17A-eGFP reporter mice were from Biocytogen. All mice were maintained under specific pathogen-free conditions. All animal experiments were performed in compliance with the guide for the care and use of laboratory animals and were approved by the institutional biomedical research ethics committee of the Shanghai Institutes for Biological Sciences (Chinese Academy of Sciences).

#### Cell culture

Naïve CD4^+^ T cells were isolated from lymph nodes or spleen of mice using a CD4^+^ CD62L^+^ T cell isolation kit II (Miltenyi Biotec) according to the manufacturer’s instruction. Naïve CD4^+^ T cells were stimulated with plate-bound anti-CD3ε mAb (5 µg/ml) in the presence of anti-CD28 mAb (2 µg/ml) in a 48-well plate under neutral conditions (10 µg/ml anti–IL-4 mAb and 10 µg/ml anti–IFN-γ mAb), IL-6 conditions (10 ng/ml IL-6, 10 µg/ml anti–IL-4 mAb, and 10 µg/ml anti–IFN-γ mAb), T_H_17 (β) condition (2 ng/ml TGF-β1, 20 ng/ml IL-6, 10 µg/ml anti–IL-4 mAb, and 10 µg/ml anti–IFN-γ mAb), T_H_17 (23) condition (20 ng/ml IL-6, 20 ng/ml IL-1β, 20 ng/ml IL-23, 10 µg/ml anti–IL-4 mAb, and 10 µg/ml anti–IFN-γ mAb).

#### In vitro PTXF treatment

Naïve CD4^+^ T cells were differentiated under pathogenic T_H_17 (23) condition for 2 days, cells were collected and pre-incubated with 500 µg/mL PTXF or H_2_O for 15 min at 37C. Total cells suspensions were then activated with 5 µg/mL plate-coated anti-mCD3, 2 µg/mL soluble anti-mCD28, 20 ng/ml IL-6, 20 ng/ml IL-1β, 20 ng/ml IL-23, 10 µg/ml anti–IL-4 mAb, and 10 µg/ml anti–IFN-γ mAb overnight at 37C.

#### Adenovirus transduction

For overexpression of GFP or c-Rel in nonpathogenic T_H_17 (β) cells, naïve CD4^+^ T cells were differentiated under nonpathogenic T_H_17 (β) condition for 2 days, then adenovirus supernatant containing GFP or c-Rel (Hanbio, Shanghai, China) was added and the cells were spun at 2000 rpm for 1.5 hr at 30C. After spin infection, the cells were cultured in T_H_17 (β) condition and harvested on day 5 for sorting of GFP^+^ transduced cells.

### Methods Details

#### Intracellular cytokine staining

Cultured or tissue isolated lymphocytes were washed and stimulated with PMA (50 ng/ml) plus ionomycin (750 ng/ml) for 4 h at 37°C. Cells were stained with anti-CD45 APC-cy7, anti-CD3ε PerCP5.5 and anti-CD4 BV510 for 30 min at 4°C. Cells were then fixed, permeabilized with Perm/Wash buffer (BD), and stained with anti–IFN-γ APC, anti-GM-CSF PE or anti–IL-17A PE for 30 min at 4°C. Cytokine profiles of CD4^+^ cells were analyzed on a FACSCalibur with FlowJo software (Tree Star).

#### Induction of EAE

IL17A-eGFP reporter mice were injected subcutaneously in the tail base with MOG35-55 peptide (200 µg/mouse, MEVGWYRSPFSRVVHLYRNGK) in complete Freund’s adjuvant. 5 min and 48 h after the injection of MOG35-55 peptide, the mice were injected intraperitoneally with pertussis toxin (200 ng/mouse; Sigma-Aldrich). Status of the mice was monitored, and disease severity was scored three times a week as follows: 0 = no clinical signs, 1 = limp tail (tail paralysis), 2 = complete loss of tail tonicity or abnormal gait, 3 = partial hind limb paralysis, 4 = complete hind limb paralysis, 5 = forelimb paralysis or moribund, 6 = death. Cells were isolated from whole brain and spinal cord at the peak of disease by Percoll gradient centrifugation (37%/70%) and subjected to flow cytometric analysis or flow cytometric sorting.

#### MicroRNA array analyses

GFP^+^ T_H_17 cells sorted from in vivo suspensions, 200-500 ileum and CNS T_H_17 cells were used for lysis, 3’ ligation, 5’ ligation, reverse transcription and preamplification using miScript single cell qPCR kit (Qiagen), cDNA was used for microRNA qPCR array analyses. The data were normalized to a U6 snRNA control.

#### RNA-seq

Naive CD4^+^ T cells from control and *miR-21*^−/−^ mice were differentiated under pathogenic T_H_17 (23) differentiation condition. Total RNA was prepared from these cells using Trizol reagent (Invitrogen). RNA-seq libraries were prepared using a TruSeq Stranded Total RNA Library Prep Kit (Illumina). For GFP^+^ T_H_17 cells sorted from in vivo suspensions, 50,000 cells were resuspended in 700 ul Trizol reagent, total RNA was prepared by miRNeasy Micro Kit (Qiagen), RNA-seq libraries were prepared using SMART-Seq v4 Ultra Low Input RNA Kit (clontech). Sequencing was performed on an Illumina HiSeq X Ten System in a 150 bp/150 bp Paired end mode.

#### ATAC-seq

ATAC-seq library preparations were performed as described(Buenrostro et al., 2013). In brief, 50,000 cells were washed in cold PBS and lysed. Transposition was performed at 37 °C for 30 min. After purification of the DNA with the MinElute PCR purification kit (Qiagen), DNA was amplified for 5 cycles. Additional PCR cycles were evaluated by real time PCR. Final product was cleaned by Ampure Beads at a 1.5× ratio. Libraries were sequenced on a HiSeq X Ten System in a 150 bp/150 bp Paired end mode.

#### Metabolomics

The metabolomics profiling was performed on gas chromatograph-time-of-flight mass spectrometry (GC-TOFMS) as previously described (Chen et al., 2014; Chen et al., 2016; Liu et al., 2019) (Metaboprofile, Shanghai, China). MetaboAnalyst was used to analyze range-scale data and provide PCA and KEGG pathway analysis of metabolites changed (www.metaboanalyst.ca/).

#### Metabolism Assays

Extracellular acidification rate (ECAR) was determined using Seahorse Xfe24 analyzer (Agilent Technologies). Briefly, T_H_17 (β), T_H_17 (23), control and *miR-21*^−/−^ T_H_17 (23) cells (0.5×10^6^/well) were plated on Cell-Tak coated Seahorse culture plate for 30 min. ECAR, a measure of glycolysis was measured under basal conditions and in response to glucose (10 mM), Oligomycin (1.0 mM), 2-deoxyglucose (2-DG) (50 mM) (Agilent seahorse XF glycolysis stress test kit).

### Quantification and Statistical Analysis

#### ATAC-seq data processing

ATAC-seq reads were trimmed using trim_galore 0.4.4_dev with the default settings then mapped to mm10 reference by bowtie2 (Langmead and Salzberg, 2012) with parameters ‘--end-to-end --no-unal -X 2000’, low quality reads (MAPQ < 10), singletons and duplicated reads were removed using samtools (Li et al., 2009). To call the differential accessible regions between samples, filtered bam files were down-sampled to the same fragments number and then processed by rgt-THOR with parameters ‘--pvalue 0.05 --binsize 100 --step 50’ (Allhoff et al., 2016). Differential peaks derived from rgt-THOR were further filtered according to their significant scores by cutting from the switch point in the rank plot. Enriched transcription factor binding motifs were searched by HOMER findMotifsGenome.pl (Heinz et al., 2010) using top 1000 differential accessible peaks centered by the peak summits and extended to +/− 100bp.

#### RNA-seq data processing

RNA-seq reads were mapped to mm10 and refGene annotation transcriptome downloaded from UCSC Genome Browser using TopHat2 (Kim et al., 2013), low quality reads (MAPQ < 10) were filtered by samtools, then gene level expression was count by htseq-count (Anders et al., 2015) using all refGenes. Differential expressed genes were called by DESeq2 (Love et al., 2014) using FDR 0.05 and foldchange ≥ 1.5.

For other quantitative data, Prism software was used for statistical analysis. Differences between groups were compared with an unpaired two-tailed t-test. A P value of less than 0.05 was considered statistically significant.

## Supporting information

Graphical abstract

## Acknowledgments

We thank Di Yu and Youcun Qian for their helpful discussions, technical expertise and/or review of this manuscript. This work is supported by grants from National Natural Science Foundation of China (31630021).

## Author Contributions

X.Y., W. J. and N.S. designed the experiments and wrote the manuscript; X.Y. did most of the experiments; L.W. helped with the mouse experiments; Z. H. helped with ATAC-seq and RNA-seq data analysis.

## Declaration of Interests

The authors declare no competing financial interests.

