## Supplemental information for "Deregulation of TGF-β1 signaling induces glycolysis by chromatin remodeling in pathogenic TH17 cells"

### Figure S1

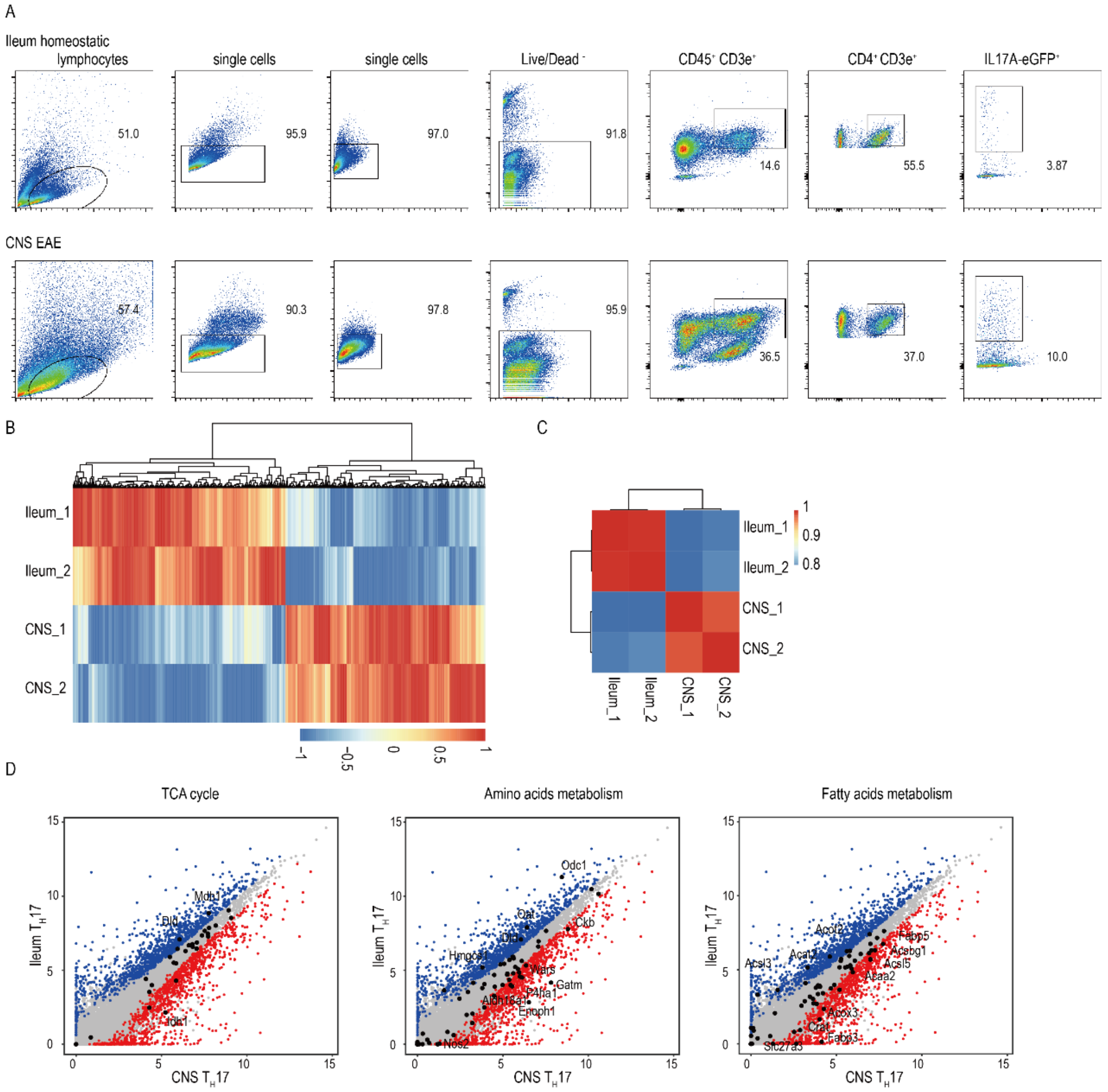


Figure S1 TH17 cells derived in vivo show discrete metabolic pathway gene expression

(A) Single cell suspensions from homeostatic ileum, EAE CNS were stained with Live/Dead, CD45, CD3e, CD4, and gated on Live/Dead - live cells for subsequent sorting for IL17A-eGFP+ TH17 cells.

(B) Heatmap shows differential expressed genes ($FDR<0.05$, $FC\geq1.5$) in $T_H17$ cells derived from homeostatic ileum and autoimmune CNS tissue.

(C) Correlative heatmap based on gene expression of indicated samples.

(D) Scatter plot shows the differential expressed genes between CNS $T_H17$ (red) and Ileum $T_H17$ (blue) cells with TCA cycle, amino acids and fatty acids pathway genes highlighted and differential genes labeled.

Figure S2

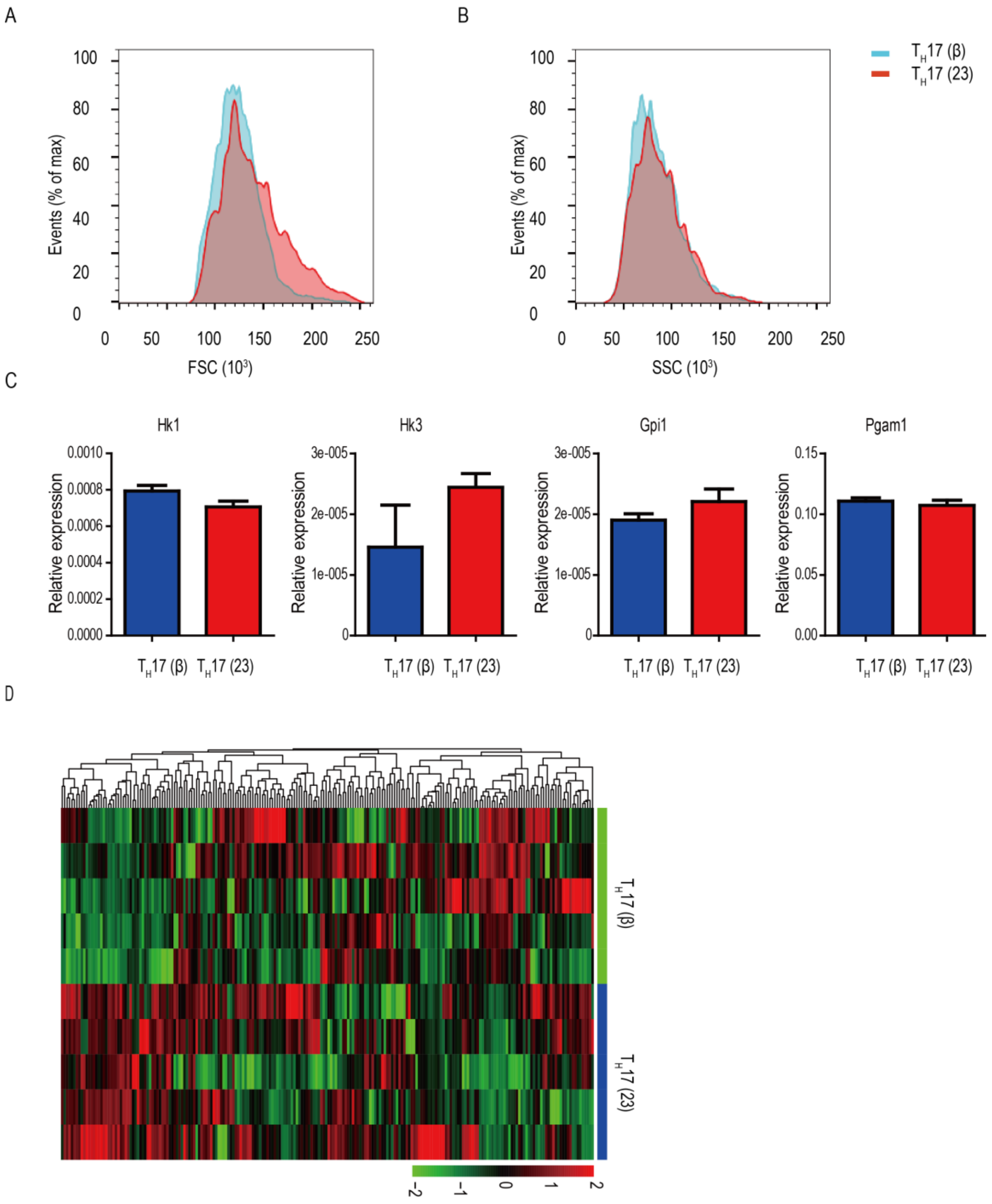


Figure S2 TH17(23) cells are metabolically different from TH17(β) cells

(A) Forward scatter plot of TH17 cells derived in vitro.

(B) Side scatter plot of TH17 cells derived in vitro.

(C) Relative expression of selected glycolytic pathway genes.

(D) The cluster heat map shows identified metabolites in $T_H17(\beta)$ and $T_H17(23)$ cells by GC-TOF/MS analysis, also see Table S1.

Figure S3

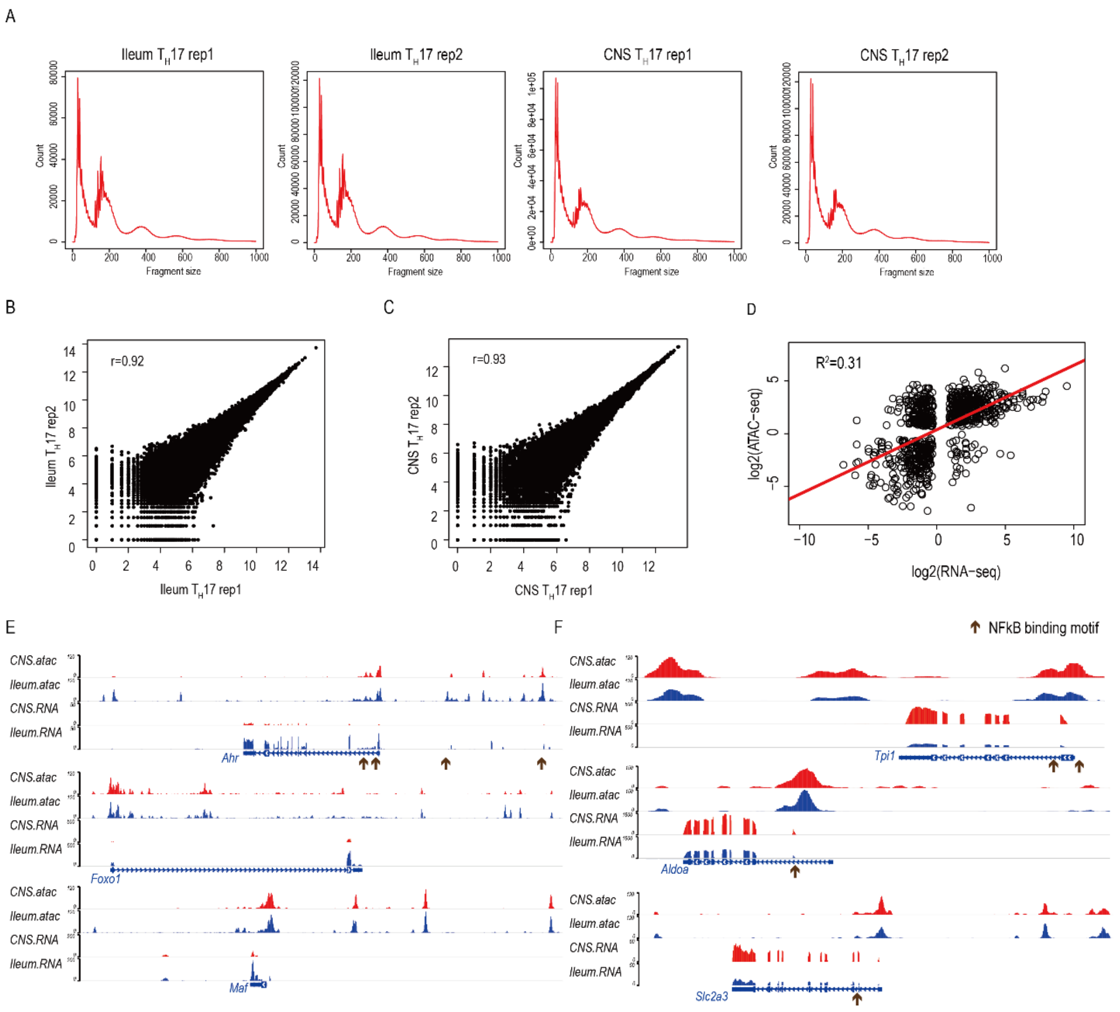


Figure S3 Quality control for ileum and CNS $T_H17$ ATAC-seq datasets

(A) Fragment size distribution for ileum and CNS $T_H17$ ATAC-seq datasets.

(B) Replication quality for ileum $T_H17$ ATAC-seq datasets.

(C) Replication quality for CNS TH17 ATAC-seq datasets.

(D) Correlation between differentially accessible regions and their associated differentially expressed genes between ileum and CNS $T_H17$ cells.

(E) Examples for differentially accessible regions and corresponding gene expression levels of $T_H17$ cell signature genes. Arrows indicate NF-kB binding motif.

(F) Examples for differentially accessible regions and corresponding gene expression levels of metabolic genes. Arrows indicate NF-kB binding motif.

Figure S4

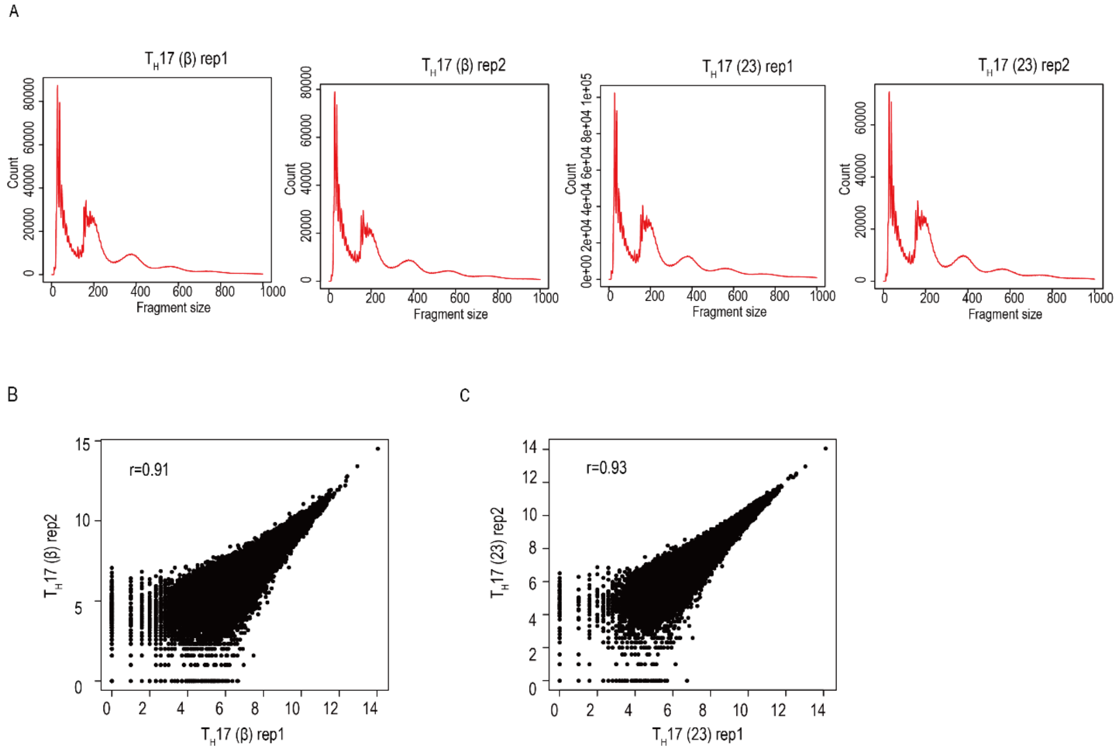


Figure S4 Quality control for in vitro derived $T_H17$ (β) and $T_H17$ (23) ATAC-seq datasets

(A) Fragment size distribution for $T_H17$ (β) and $T_H17$ (23) ATAC-seq datasets.

(B) Replication quality for $T_H17$ (β) ATAC-seq datasets.

(C) Replication quality for $T_H17$ (23) ATAC-seq datasets.

Figure S5

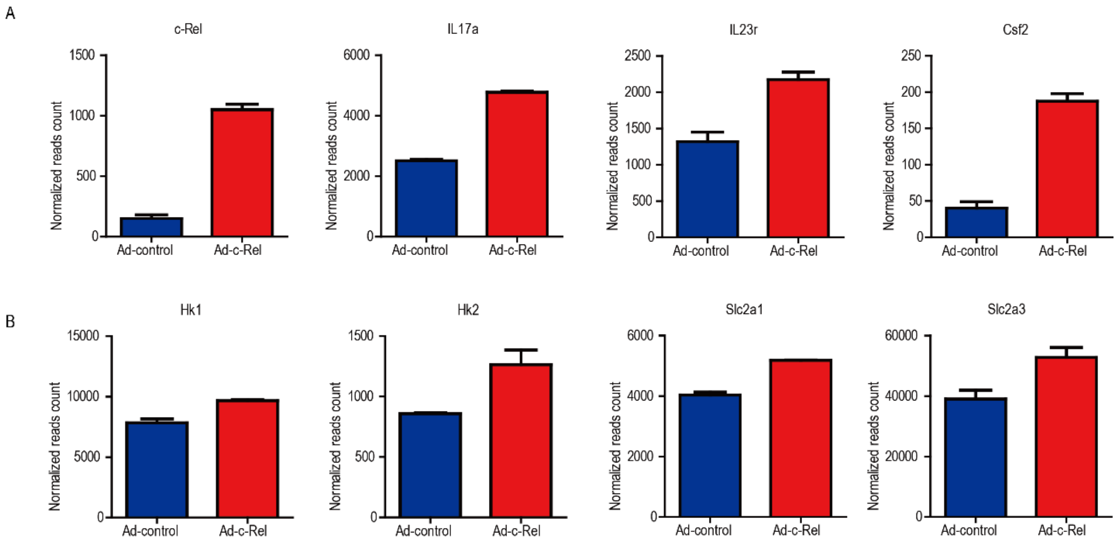


Figure S5 c-Rel overexpression in $T_H17$ (β) cells promotes pathogenic $T_H17$ cell signature gene and glycolytic gene expression

(A) Normalized reads count of selected pathogenic signature genes.

(B) Normalized reads count of selected glycolytic genes.

Figure S6

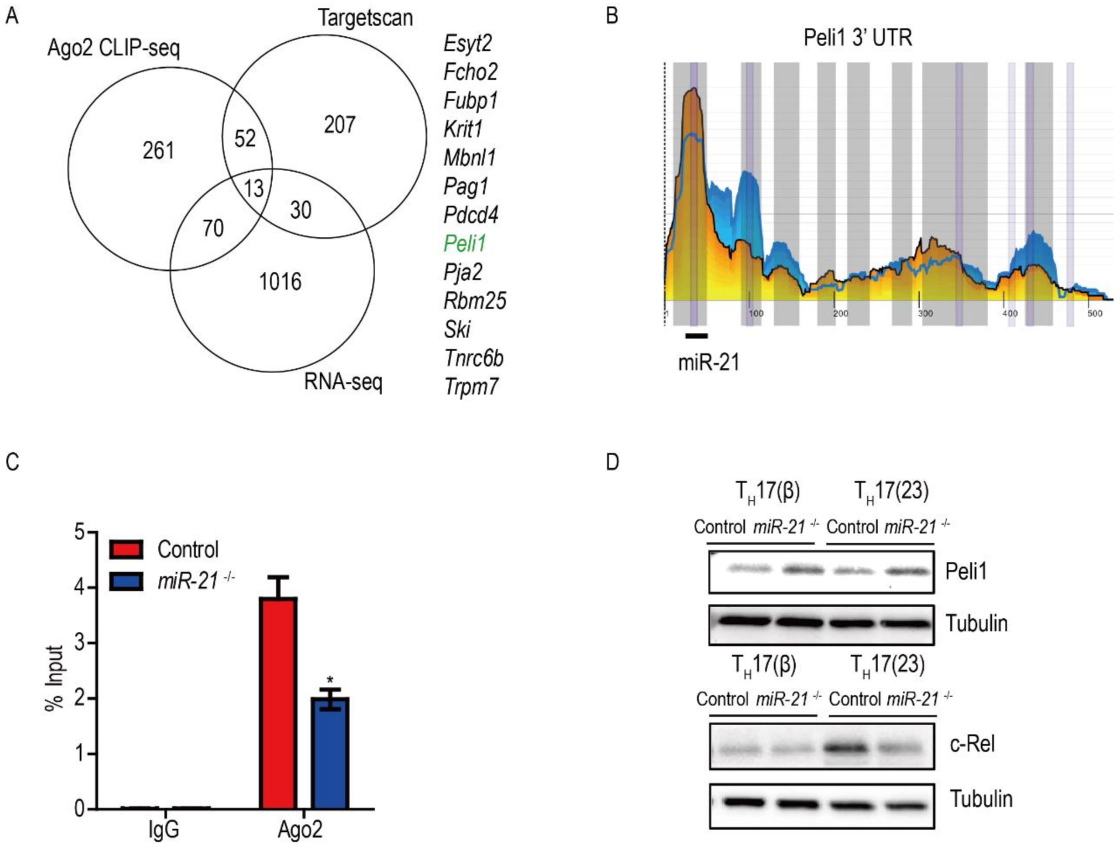


Figure S6 MiR-21 promotes glucose metabolism of pathogenic $T_H17$ cells

(A) Comparison of gene list from RNA-seq, Targetscan and CLIP-seq data, showing an overlap of 13 potential miR-21 target genes.

(B) The 3' UTR of Peli1 in activated $CD4^+$ T cells from Ago2 HITS-CLIP data (Loeb et al., 2012).

(C) RT-PCR analysis of enriched Peli1 mRNA from Ago2 immunoprecipitated total RNA from in vitro differentiated control and *miR-21*$^{-/-}$ $T_H17(23)$ cells.

(D) Western blot analysis of c-Rel and Peli1 from $T_H17$ cells differentiated under indicated condition.

*$P < 0.05$ (unpaired t-test). Data are from one experiment representative of two independent experiments (mean and s. d. in C, D).

Table S1 The Significantly different metabolites in our metabolomics data

| Table S1 Differential metabolites between the group using univariate statistical analysis | | | | | |
|---|---|---|---|---|---|
| Class | Name | HMDBID | KeggID | p | FC |
| Alkylamines | Ratio of Spermidine/Putrescine | HMDB01257/HMDB01414 | C00315/C00134 | 1.40E-03 | 0.6 |
| | Putrescine | HMDB01414 | C00134 | 2.80E-03 | 1.8 |
| Amino Acid | Ratio of L-Glutamic acid/Pyroglutamic acid | HMDB00148/HMDB00267 | C00025/C01879 | 2.70E-06 | 0.5 |
| | L-Alloisoleucine | HMDB00557 | NA | 2.90E-05 | 1.7 |
| | L-Tyrosine | HMDB00158 | C00082 | 3.20E-04 | 1.7 |
| | L-Lysine | HMDB00182 | C00047 | 6.10E-04 | 1.5 |
| | L-Valine | HMDB00883 | C00183 | 1.40E-03 | 1.6 |
| | Ratio of L-Tyrosine/L-Phenylalanine | HMDB00158/HMDB00159 | C00082/C00079 | 1.90E-03 | 1.2 |
| | L-Phenylalanine | HMDB00159 | C00079 | 3.30E-03 | 1.5 |
| | L-Glutamic acid | HMDB00148 | C00025 | 4.30E-03 | 0.6 |
| | Creatinine | HMDB00562 | C00791 | 5.80E-03 | 0.3 |
| | Ratio of Putrescine/Ornithine | HMDB01414/HMDB00214 | C00134/C00077 | 6.20E-03 | 1.6 |
| | Ratio of L-Asparagine/L-Aspartic acid | HMDB00168/HMDB00191 | C00152/C00049 | 7.50E-03 | 1.5 |
| | Ornithine | HMDB00214 | C00077 | 9.60E-03 | 1.3 |
| | Pyroglutamic acid | HMDB00267 | C01879 | 1.30E-02 | 1.3 |
| | L-Threonine | HMDB00167 | C00188 | 1.80E-02 | 1.4 |
| | L-Serine | HMDB00187 | C00065 | 3.70E-02 | 1.4 |
| | L-Alanine | HMDB00161 | C00041 | 4.30E-02 | 1.5 |
| | L-Isoleucine | HMDB00172 | C00407 | 0.057 | 1.5 |
| | Gamma-Aminobutyric acid | HMDB00112 | C00334 | 0.06 | 0.7 |
| | L-Asparagine | HMDB00168 | C00152 | 0.071 | 1.2 |
| | L-Proline | HMDB00162 | C00148 | 0.083 | 1.1 |
| Carbohydrates | D-Glucose | HMDB00122 | C00031 | 4.60E-05 | 1.2 |
| | Gluconic acid | HMDB00625 | C00257 | 2.10E-04 | 0.4 |
| | Mannitol | HMDB00765 | C00392 | 3.50E-02 | 1.2 |
| | D-Threitol | HMDB04136 | C16884 | 0.071 | 1.3 |
| Fatty Acids | Palmitoleic acid | HMDB03229 | C08362 | 0.096 | 0.7 |
| Lipids | Cholesterol | HMDB00067 | C00187 | 4.90E-03 | 0.9 |
| | Glycerol 3-phosphate | HMDB00126 | C00093 | 1.20E-02 | 1.2 |
| | O-Phosphoethanolamine | HMDB00224 | C00346 | 2.00E-02 | 0.9 |

| | | | | | |
|---|---|---|---|---|---|
| Nucleotide | Ratio of Adenine/Adenosine | HMDB00034/HMDB00050 | C00147/C00212 | **1.10E-04** | **6.3** |
| | Adenosine | HMDB00050 | C00212 | **2.60E-04** | **0.3** |
| | Inosine | HMDB00195 | C00294 | **5.00E-04** | **0.3** |
| | Guanosine | HMDB00133 | C00387 | **5.80E-04** | **0.4** |
| | Uridine | HMDB00296 | C00299 | **1.20E-03** | **0.5** |
| | Ratio of Inosine/Adenosine | HMDB00195/HMDB00050 | C00294/C00212 | **3.40E-03** | **1.1** |
| | Adenine | HMDB00034 | C00147 | **1.50E-02** | **1.5** |
| Organic Acids | Hydroxyphenyllactic acid | HMDB00755 | C03672 | **9.30E-04** | **1.2** |
| | Malic acid | HMDB00744 | C00711 | **1.20E-02** | **1.4** |
| | Taurine | HMDB00251 | C00245 | **1.80E-02** | **0.9** |
| | Fumaric acid | HMDB00134 | C00122 | **0.08** | **1.3** |
| | Pyruvic acid | HMDB00243 | C00022 | **0.088** | **1.2** |

Table S2 Primers for qPCR

| Gene name | Sequence |
|---|---|
| *miR-21* | cgtagcttatcagactgatgttga |
| *miR-155* | ttaatgctaattgtgataggggt |
| *miR-146a* | tgagaactgaattccatgggtt |
| *miR-30e* | tgtaaacatccttgactggaag |
| *miR-210* | cgtgtgacagcggctga |
| *miR-183* | tatggcactggtagaattcact |
| *miR-182* | gcaatggtagaactcacaccg |
| *miR-22* | aagctgccagttgaagaactgt |
| *miR-27a* | cacagtggctaagttccgc |
| *miR-24* | gctcagttcagcaggaacag |
| *miR-148a* | tcagtgcactacagaactttgt |
| *miR-191* | ggaatcccaaaagcagctg |
| *miR-148b* | tcagtgcatcacagaactttgt |
| *miR-186* | caaagaattctccttttgggct |
| *miR-17* | caaagtgcttacagtgcaggtag |
| *miR-98* | tgaggtagtaagttgtattgtt |
| *U6* | agattagcatggcccctg |
| *Glut1* f | GAGTGCAGGGAGGAGAGGG |
| *Glut1* r | GTGTCCGTGTCTTCAGCAGTT |
| *Hk2* f | TGCTAGGTTGACAGCTCTCTCT |
| *Hk2* r | ACAAAACGCTCACTAGACCGA |
| *Peli1* f | CCAGCAGACCAGCCTTAACT |
| *Peli1* r | GTCCTCACTTTCATCCGCTAA |
| *actb* f | GGCTGTATTCCCCTCCATCG |
| *actb* r | CCAGTTGGTAACAATGCCATGT |
