## Supplementary figures and images for "Deregulation of TGF-β1 signaling induces glycolysis by chromatin remodeling in pathogenic TH17 cells"

### Graphical abstract

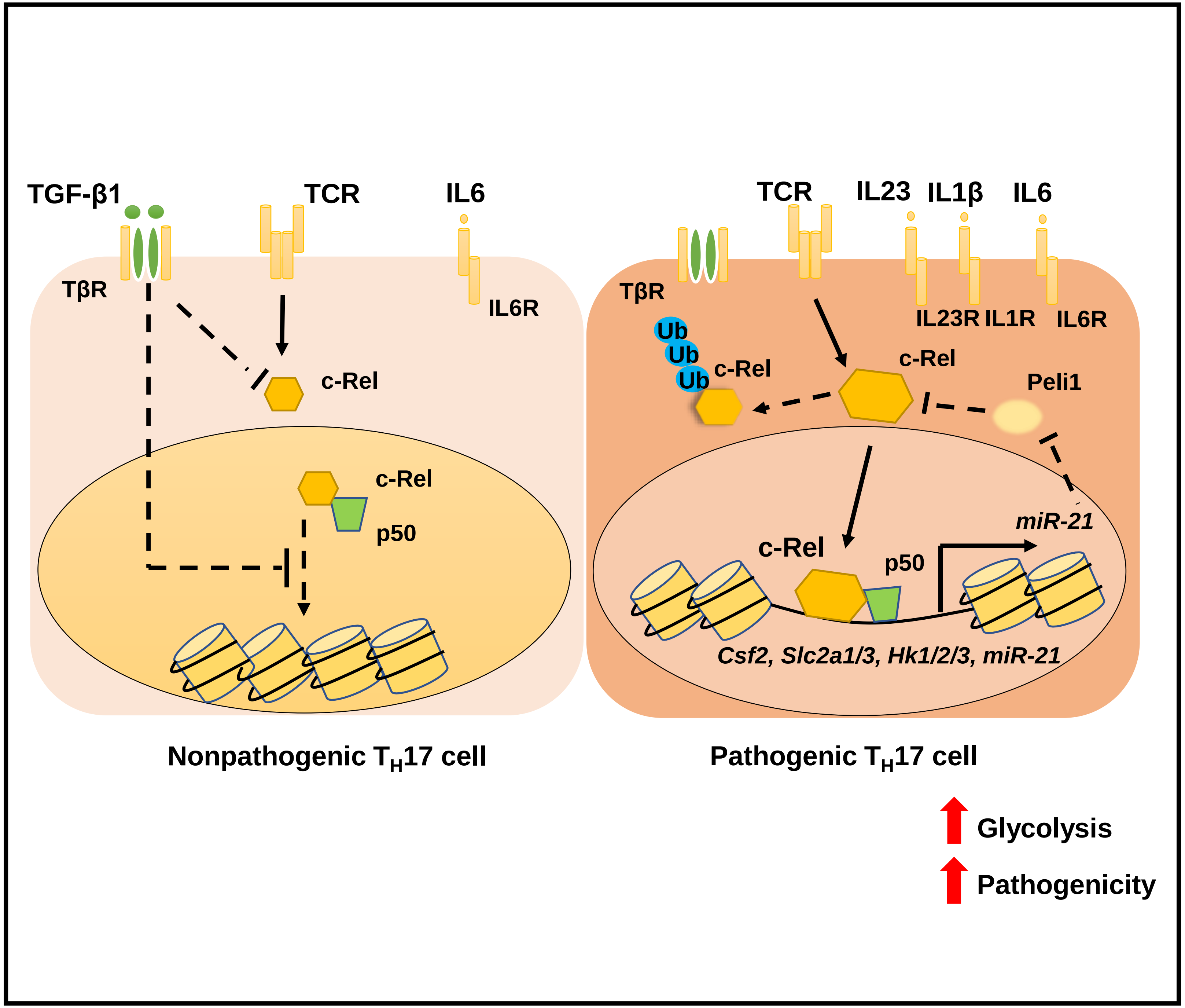
